# miR858a-encoded peptide, miPEP858a, interacts with miR858a promoter and requires C-terminus for the associated functions

**DOI:** 10.1101/2024.02.16.580742

**Authors:** Himanshi Gautam, Ashish Sharma, Anwesha Anyatama, Prabodh Kumar Trivedi

**Affiliations:** CSIR-National Botanical Research Institute, Council of Scientific and Industrial Research (CSIR-NBRI), Rana Pratap Marg, Lucknow-226001, India; Academy of Scientific and Innovative Research (AcSIR), Ghaziabad-201002, India; CSIR-Central Institute of Medicinal and Aromatic Plants (CSIR-CIMAP) P.O. CIMAP, Near Kukrail Picnic Spot, Lucknow-226 015, India

**Keywords:** Arabidopsis, miRNA, miPEP, C-terminal peptide, flavonoid, gene

## Abstract

MicroRNAs (miRNAs) play a crucial role in regulating gene expression as processed products of primary miRNAs (pri-miRNAs). Recent studies suggest that certain pri-miRNAs encode miRNA-encoded Peptides (miPEPs), influencing miRNA function. Despite this emerging insight, molecular mechanisms and the specific domain or essential amino acid residues required for the functionality of these miPEPs have yet to be explored. Here, we report that the pri-miR858a-encoded peptide, miPEP858a, interacts with the promoter region of the MIR858 gene and its C-terminal region consisting of 14 amino acid residues is indispensable for its functionality. DNA-protein interaction through Yeast-one-hybrid and ChIP-qPCR analysis revealed that miPEP858 interacts with specific region in the MIR858 promoter. An exogenous application of a truncated peptide, the C-terminal region, resulted in enhanced expression and related phenotypic effects, similar to complete miPEP858. Furthermore, C-terminal region of miPEP858a complemented mutant plants lacking miPEP858, highlighting their regulatory role in miR858a expression and associated target genes, impacting flavonoid biosynthesis and plant development. These findings suggest that full-length miPEP858a may not be essential for its functionality. Identifying functional domains of other peptides offers promise, potentially reducing synthesis costs for application in crop plants and avoiding laborious biotechnological approaches to enhance agronomic traits.

## Introduction

The increasing world population is challenging and always concerned with food safety and security. Due to poor yield, drastic changes in environmental conditions affect crop production, and food security is debatable nowadays. Thus, modern science, especially biotechnology, paving the way for this emerging issue to ensure food security and higher agricultural produce with lesser yield losses (Moore et al., 2021, Chakraborty et al., 2011, Li et al., 2022, Zhang et al., 2020). It has been estimated about one-third of agricultural produce depends upon the use of pesticides, impacting human health and the rest is affected by climate change (Tudi et al., 2021). Thus, crop improvement with novel, easy, and safe methods is the utmost requirement to meet the demands of the population (Ormancey et al., 2023, Ormancey et al., 2021). Advancements in computational approaches, peptidomics and transcriptomics allow us to identify the various regulatory peptides in plants that can be potent tools to improve agronomic traits (Tavormina et al., 2015, Fabre et al., 2021).

It has been well demonstrated that primary transcripts of miRNA encode regulatory peptides called microRNA-encoded peptides (Lauressehues et al., 2015). In soybean, it has been observed that miPEP172c increases the expression of miR172c, thus ultimately increasing nodule number (Couzigou et al., 2016). Functional characterization of small peptide encoded by miR171d, vvi-miPEP171d, promotes the adventitious root development by activating the expression of vvi-miR171d (Chen et al., 2020). Exogenous application of miPEP164c is also known to be involved in inhibiting the proanthocyanidin content through acting on its pathway and facilitating the accumulation of anthocyanin biosynthesis (Vale et al., 2021). Another fully characterized miR858a encoded peptide, miPEP858, regulates the expression of miR858 and target genes, resulting in modulated levels of flavonoids due to changes in the expression of genes associated with phenylpropanoid pathway, auxin signalling and regulated by light dependent transcription factor, HY5 (Sharma et al., 2020, Sharma et al., 2022). Another report suggests the involvement of miR408, which targets a glutathione S-transferase GSTU25 and exogenous application of miR408-encoded-peptide, miPEP408, plays a role in sulfur assimilation and to detoxify the environmental pollutant (Kumar et et al., 2023). The external application of synthetic Vvi-miPEP172b and Vvi-miPEP3635b to the grape plantlets enhances cold tolerance compared to control conditions (Chen et al., 2022). Till now, little is known about the mechanism of action of miPEPs; however, the molecular basis of miPEP specificity has been deciphered that suggests the miPEP activity relies on its own miORF (Lauressergues et al., 2022). Altogether, these studies suggest the potential role of these miPEPs in enhancing desired traits in several crop plants. In addition, previous studies on miPEPs confined to identification, characterizing their function and effects of exogeneous treatment for enhancing the phenotype and other associated functions.

Major bottlenecks regarding the synthesis of peptides, even in smaller concentration, is quite expensive and difficult to adopt for agricultural produce (Feng et al., 2023). Majority of small signaling peptides undergo C-terminal processing occurs to produce mature signals (Mastubayashinet al., 2011, Mastubayashinet al., 2014). A study showed that AtPEP3, a peptide that showed its potential to increase salinity stress tolerance, has minimal functional fragment that has the potential to enhance salinity stress tolerance (Nakaminami et al., 2018). Our earlier study showed that exogenous application of miPEP858 affects flavonoid biosynthesis in addition to plant growth and development. Therefore, we hypothesised whether a functional minimal fragment in miPEP858 is cost-effective and functions similarly to miPEP858.

We report that the miPEP858 interacts with the promoter of MIR858 gene and C-terminus of miPEP858 is essentially required for regulating the expression of miR858 and target gene expression. Previous studies suggested miR858-encoded peptide, miPEP858 is involved in regulating the expression of miR858 target (R2R3-MYB (MYB111, MYB11, MYB12) genes and associated gene functions (Sharma et al., 2020, Sharma et al., 2016, Tirumalainet al., 2019, Piya et al., 2017). We divided full-length miPEP858 into different-sized fragments and used these individually to decipher the function. Exogenous application of all the truncated peptides suggests the C-terminal amino acids are the most responsive out of all the truncated peptides. Promoter reporter analysis through exogenous application of all truncated peptides revealed the increased GUS expression with C-terminus peptides truncated peptide similar to full-length miPEP858a. The complementation of *miPEP858^CR^* lines with truncated peptide also led to the restoration of growth-related phenotype. Moreover, we also identified the effect of truncated peptides at the CHS and MYB12 protein levels due to an increase in the expression of mature and pre-miR858 and subsequently decreased the protein accumulation of target genes. This study will increase the focus on identifying the miPEPs in other crops and their minimum functional fragments to pave the way for utilising them in agriculture to enhance traits in several crops.

## Results

### miPEP858a interacts on its promoter and regulate expression of miR858a

Several reports have been published on the characterization of plant miPEPs; however, the molecular mechanism underlying miPEP action remains elusive. The application of exogenous miPEP to promoter-reporter lines has been shown to enhance reporter expression (Lauressegues et al., 2015, Sharma et al., 2020, Kumar et al., 2023, Lauressegues et al., 2022). Notably, recent evidence in animals indicates the regulation of miR-31 expression through its encoded peptide, miPEP31 (Zhou et al., 2022). MiPEP31 functions as a transcriptional repressor, negatively influencing miR-31 expression by binding to its promoter region. In Arabidopsis, promoter: reporter studies of miPEP858a also support enhanced GUS gene activity and expression upon peptide supplementation. This implies that miPEP858a can bind to the promoter region of miR858a, thereby regulating its promoter activity. To unravel the molecular mechanism of miPEP function, various approaches were employed, including the exploration of possible mechanisms such as the modeling of 3D structures and its binding on the promoter. Bioinformatics analysis suggests three putative binding sites (R1, R2, and R3) for miPEP858a on the miR858a promoter (Figure 1A, Supplemental Figure 1).

**Figure 1.**
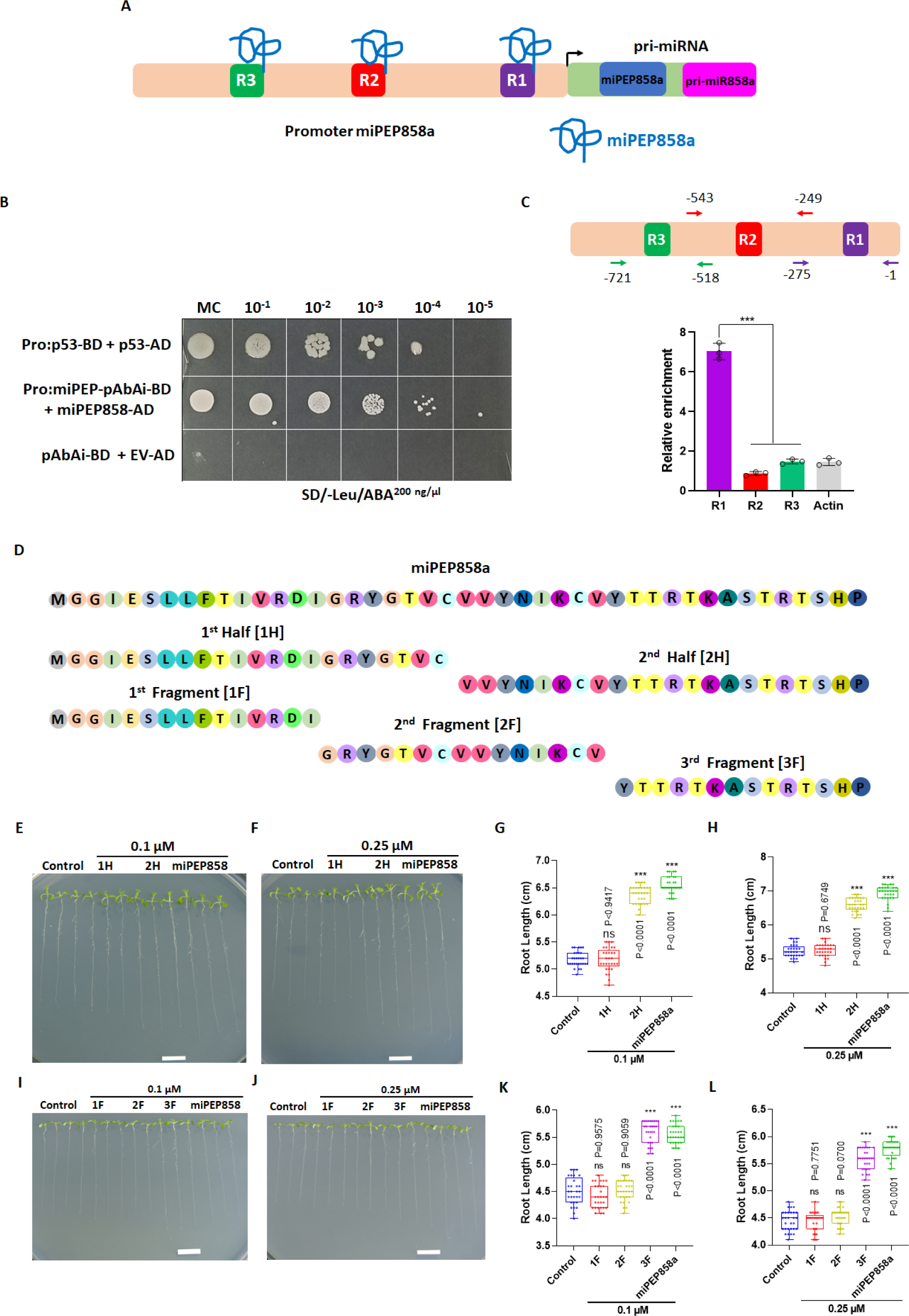
Interaction of miPEP858a with its own promoter and identification of functional truncated peptides. **(A)** Schematic diagram showing three putative binding site (R1, R2 and R3) of miPEP858a on promoter of miPEP858a. **(B)** Yeast one hybrid assay showing interaction between miPEP858a and miPEP858a promoter. **(C)** Promoter region showing primer location for each putative binding site used for Chip assay. Graph showing relative enrichment of the fragment. **(D)** Schematic representation of parent peptide encoded by pri-miR858a (miPEP858a: 44 amino acids residues) and its truncated sequences: 1st Half (1H-22 amino acids) and 2nd Half (2H-22 amino acids); 1^st^ Fragment (1F-15 amino acids), 2^nd^ Fragment (2F-15 amino acids) and 3^rd^ Fragment (3F-14 amino acids). **(E)** Representative image of ten-day-old WT seedlings grown on half-strength MS medium supplemented with water (control) or 0.1 µM of truncated peptide 1H, 2H and miPEP858a. Scale bar, 1 cm. **(F)** Representative image of ten-day-old WT seedlings grown on half-strength MS medium supplemented with water (control) or 0.25 µM of truncated peptide 1H, 2H and miPEP858a. Scale bar, 1 cm. **(G)** Root lengths of ten-day-old WT seedlings grown on half-strength MS medium supplemented with water (control) or 0.1 µM of truncated peptide 1H, 2H and miPEP858a. n= 30 independent seedlings (small open circles). **(H)** Root lengths of ten-day-old WT seedlings grown on half-strength MS medium supplemented with water (control) or 0.25 µM of truncated peptide 1H, 2H and miPEP858a. n= 30 independent seedlings (small open circles). **(I)** Representative image of ten-day-old WT seedlings grown on half-strength MS medium supplemented with water (control) or 0.1 µM of truncated peptide 1F, 2F,3F and miPEP858a. Scale bar, 1 cm. **(J)** Representative image of ten-day-old WT seedlings grown on half-strength MS medium supplemented with water (control) or 0.25 µM of truncated peptide 1F, 2F,3F and miPEP858a. Scale bar, 1 cm. **(K)** Root lengths of ten-day-old WT seedlings grown on half-strength MS medium supplemented with water (control) or 0.1 µM of truncated peptide 1F, 2F, 3F and miPEP858a. n= 30 independent seedlings (small open circles). **(L)** Root lengths of ten-day-old WT seedlings grown on half-strength MS medium supplemented with water (control) or 0.25 µM of truncated peptide 1F, 2F, 3F and miPEP858a. n= 30 independent seedlings (small open circles).

To unravel the in-vivo molecular mechanism associated with miPEP858a, a Yeast-One Hybrid (Y1H) assay was conducted, examining the interaction between the miPEP858a promoter (used as bait) and the miPEP858 protein (prey). The Y1H results indicated a robust interaction between the miPEP858a promoter and miPEP858a (Figure 1B). To further corroborate this interaction, a Chromatin Immunoprecipitation (ChIP) assay was executed. Analysis of relative enrichment via ChIP-qPCR revealed enhanced levels of DNA fragments containing the R1 binding site compared to the other two putative binding sites (Figure 1C). Collectively, these findings unequivocally demonstrate that miPEP858a possesses the potential to bind to its promoter, thereby regulating the expression of miR858a.

### Identification of functional truncated peptides

The primary transcript of miRNA has been found to encode microRNA encoded peptides (miPEPs), which enhance the transcription of their corresponding miRNAs (Lauressegues et al., 2015, Gautam et al., 2023). To identify functional domains within miPEPs or their potential as full-length peptides, we focused on miPEP858, known to modulate the expression of miR858 and its target genes. Two truncated peptides, 1H (N-terminal 22 amino acids) and 2H (remaining 22 amino acids), were designed (Figure 1D). Exogenous application of miPEP858a enhances primary root length (Sharma et al., 2020), leading us to hypothesize that these truncated peptides would exhibit similar functionality if active. Root length data revealed that 1H had no significant effect, while 2H significantly increased root length in a concentration-dependent manner compared to control seedlings (Figure 1E-H).

Additionally, smaller truncated peptides were designed by fragmenting the full-length peptide into three segments: 1F (N-terminus, 15 amino acids), 2F (middle region, 15 amino acids), and 3F (C-terminus, 14 amino acids) (Figure 1D). Growing WT seeds on media supplemented with these peptides at two concentrations indicated that 3F had the potential to enhance root length, similar to miPEP858a, while 1F and 2F showed no significant changes in root length (Figure 1I-L). In summary, these results highlight the potential of 2H and 3F peptides, containing C-terminal amino acids, to modulate root length similarly to miPEP858.

### Modulation of miR858 and associated genes expression by truncated peptides

Previously, full-length miPEP858 was recognized for enhancing miR858 expression and subsequently downregulating its target and associated genes (Lauressegues et al., 2015). Building on our initial findings, we hypothesized that if 2H and 3F can enhance root length, they may similarly impact miR858 expression. Expression analysis revealed a significant increase in mature miR858a expression when treated with 2H and 3F, while other truncated peptides showed no significant difference compared to control seedlings (Figure 2A). Additionally, analysis of pre-miR858, target, and associated genes demonstrated elevated pre-miR858 expression and significant downregulation of target genes (Figure 2B). To validate these results, we examined MYB12 and CHS protein accumulation using specific antibodies. Functional truncated peptides (2H and 3F) led to a significant decrease in protein levels, akin to full-length miPEP858, compared to other truncated peptides (Figure 2C). Since miR858 targets genes in the phenylpropanoid pathway (Sharma et al., 2016), our results indicated the downregulation of genes encoding MYB12 and CHS upon supplementation with 2H and 3F truncated peptides. Additionally, we analyzed the expression of other pathway genes with all truncated peptides. The results suggested significant downregulation with 2H and 3F, and no significant modulation with 1H, 1F, and 2F truncated peptides (Supplemental Figure 2). In summary, these findings highlight the crucial role of the minimal region 2H and 3F truncated peptides in downregulating the expression of genes involved in the phenylpropanoid pathway.

**Figure 2.**
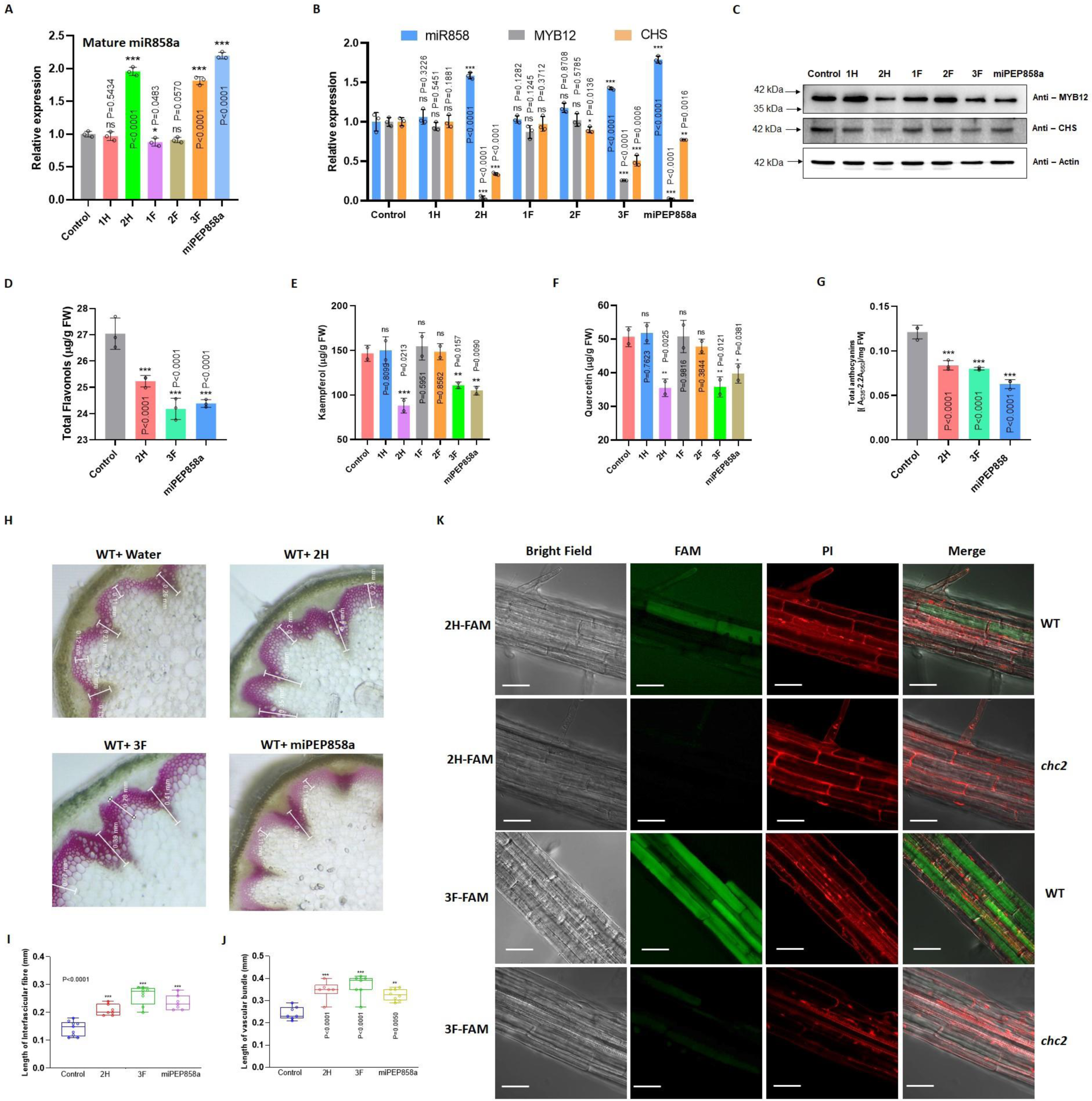
Truncated peptides modulate the expression, accumulation of secondary metabolites and follow clathrin-mediated endocytosis. **(A)** Quantification of mature miR858a using TaqMan probe assay in five-day-old WT seedlings grown on half-strength MS medium supplemented with water (control) and 0.25 µM of 1H, 2H, 1F, 2F, 3F and miPEP858a. **(B)** Quantification of miR858a and its target genes in five-day-old WT seedlings grown on half-strength MS medium supplemented with water (control) and 0.25 µM of 1H, 2H, 1F, 2F, 3F and miPEP858a. **(C)** Western blot analysis of MYB12 and CHS protein in five-day-old WT seedlings grown on half-strength MS medium supplemented with water (control) and 0.25 µM of 1H, 2H, 1F, 2F, 3F and miPEP858a. Actin was used as the loading control. **(D)** Quantification of total flavonol in ten-day-old WT seedlings grown on half-strength MS medium supplemented with water (control) and 0.25 µM of functional truncated peptides 2H, 3F and miPEP858a. FW, fresh weight. **(E)** Quantification of kaempferol content in ten-day-old WT seedlings grown on half-strength MS medium supplemented with water (control) and 0.25 µM of 1H, 2H, 1F, 2F,3F and miPEP858a. FW, fresh weight. **(F)** Quantification of quercetin content in ten-day-old WT seedlings grown on half-strength MS medium supplemented with water (control) and 0.25 µM of 1H, 2H, 1F, 2F,3F and miPEP858a. FW, fresh weight. **(G)** Quantification of anthocyanin in ten-day-old WT seedlings supplemented with water (control) and 0.25 µM of 2H, 3F and miPEP858a. FW, fresh weight. **(H)** Representative images of transverse sections of stems of 35-day-old WT plants supplemented with water (control) and 0.25 µM of 2H, 3F and miPEP858a stained with phloroglucinol showing changes in lignin content. The experiment was repeated three times with n= 5 biologically independent replicates, with similar results. **(I)** Lengths of interfascicular fibres of 35-day-old stems of WT plants supplemented with water (control) and 0.25 µM of 2H, 3F and miPEP858a (n= 15). **(J)** Lengths of vascular bundles of 35-day-old stems of WT plants supplemented with water (control) and 0.25 µM of 2H, 3F and miPEP858a (n= 15). **(K)** Confocal images showing the uptake of FAM-labeled 2H and 3F peptides (5-FAM 2H and 5-FAM 3F) and PI in WT and *chc2* roots after incubation for 12 h. Scale bars, 20 µM. Confocal images are representative of three independent experiments, n = 10 seedlings.

### Functional truncated peptides modulate metabolite content

Exogenous application of C-terminal truncated peptides enhanced the expression of key genes in the phenylpropanoid pathway, prompting an analysis of metabolite accumulation. Total flavonol content analysis indicated a significant decrease in seedlings treated with functional truncated peptides (2H, 3F) compared to those supplemented with non-functional truncated peptides (Figure 2D). In-depth HPLC analysis quantifying kaempferol and quercetin levels revealed a significant decrease in seedlings treated with functional truncated peptides compared to the control (Figure 2E and 2F, Supplemental Figure 3). Total anthocyanin analysis suggested reduced accumulation in seedlings grown on media supplemented with the C-terminal peptide (Figure 2G). In line with our previous study indicating increased flavonol and anthocyanin levels at the expense of lignin production, we analyzed lignin levels under different growth conditions. Cross-section staining of WT plants treated twice with truncated peptide confirmed significantly enhanced lignification in interfascicular and vascular tissues (Figure 2H). Additionally, the length of interfascicular fibers and vascular bundles increased when supplemented with 2H and 3F truncated peptides (Figure 2I and 2J). These results underscore the potential of the functional region (2H and 3F) of miPEP858 to significantly alter metabolite levels.

### Clathrin-mediated endocytosis is required for truncated peptides for internalization

Functional truncated peptides (2H and 3F) have demonstrated significant effects on key genes in the phenylpropanoid pathway, influencing plant growth and development. Previous studies have indicated the necessity of clathrin-mediated internalization for miPEP entry into plant cells (Sharma et al., 2020, Ormancey et al., 2020, Badola et al., 2022). To investigate the entry mode of truncated peptides, FAM-labeled 2H and 3F peptides were synthesized. WT and clathrin mutant (*chc1, chc1.2, chc2, chc2.2*) seedlings were incubated with these peptides, and microscopic observation revealed fluorescence within plant cells in WT seedlings, while clathrin mutants showed no internal fluorescence (Figure 2K, Supplemental Figure 4 and 5). Furthermore, clathrin mutants were grown on media with and without truncated peptides. Root length analysis indicated no effect on the root length of clathrin mutants with the exogenous application of C-terminal peptide and other non-functional truncated peptides (Supplemental Figure 6 and 7). In conclusion, our results confirm clathrin-mediated endocytosis of FAM-labeled 2H and 3F peptides, facilitating internalization into plant cells, similar to previous findings with full-length miPEP858a.

### Complementation of *miPEP858^CR^* lines by truncated peptides

The exogenous application of miPEP858 is known to complement the phenotype and molecular function of *miPEP858^CR^* lines (Sharma et al., 2020). To study whether truncated peptides can restore the phenotype of *miPEP858^CR^* seedlings, *miPEP858^CR^*and WT seedlings were grown on media supplemented with non-functional and functional truncated peptides. Results suggested the increase in root length and overall development of the *miPEP858^CR^* seedling supplemented with functional truncated peptides (2H and 3F truncated peptides) along with full-length miPEP858 as a positive control (Figure 3A-D). To further determine that functional truncated peptides affect *miPEP858^CR^* seedlings via miR858 associated pathway, miR*858^CR^* seedlings were grown on media supplemented with functional truncated peptides. No modulation in the root length of *miR858^CR^* lines was observed when supplemented with truncated peptides along with miPEP858 (Supplemental Figure 8). This suggests that truncated peptides modulate the phenotype through miR858 associated pathway.

**Figure 3.**
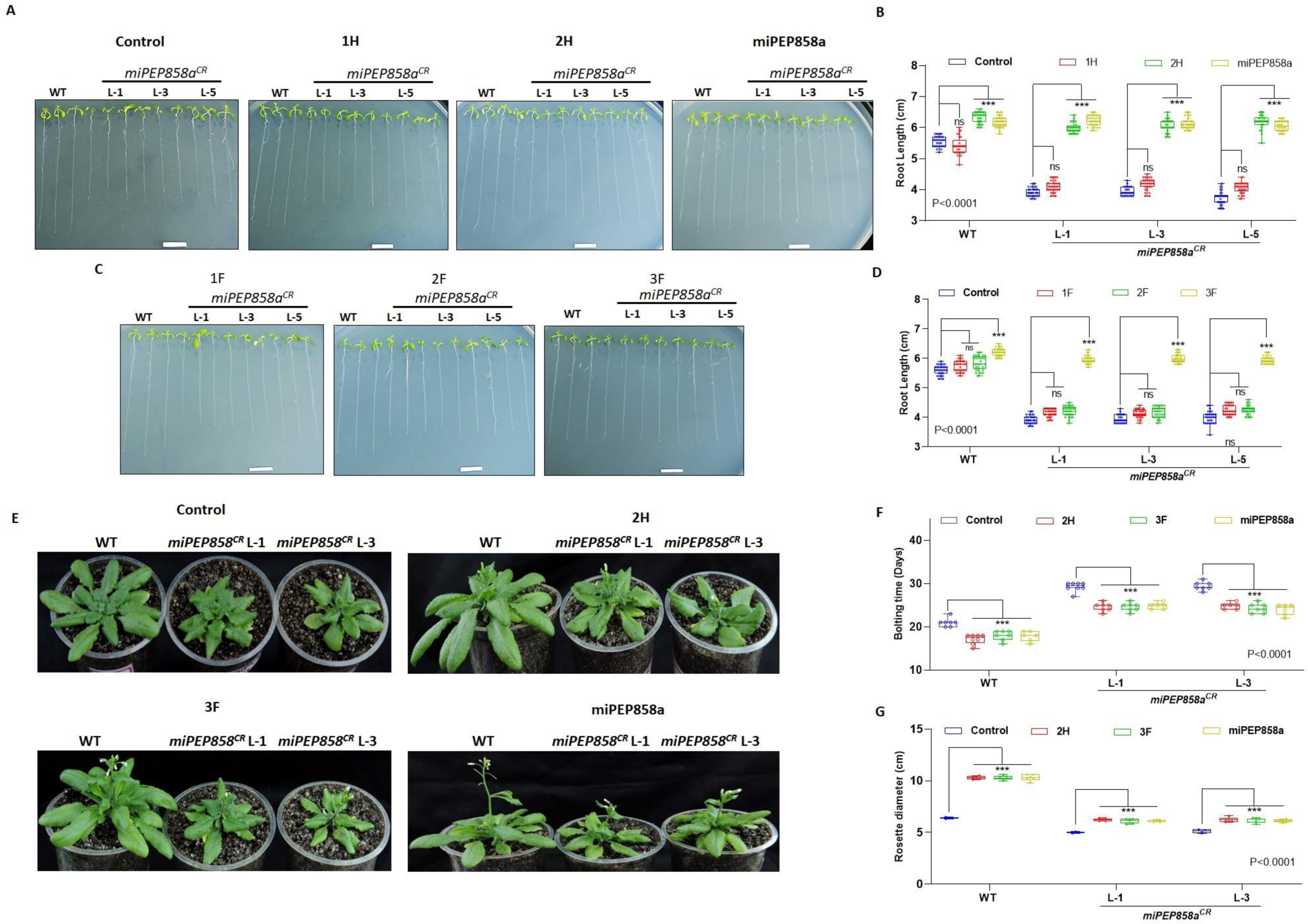
Complementation of *miPEP858^CR^* phenotype by functional truncated peptide via miR858-associated pathway. **(A)** Representative image of ten-day-old seedlings of WT and miPEP858^CR^ lines (L-1, L-3, L-5) grown on half-strength MS medium supplemented with water (control) and 0.25 µM 1H, 2H and miPEP858a. Scale bar, 1 cm. **(B)** Root lengths of ten-day-old seedlings of WT and *miPEP858^CR^* lines (L-1, L-3, L-5) grown on half-strength MS medium supplemented with water (control) and 0.25 µM 1H, 2H and miPEP858a (n= 30 independent seedlings, indicated by the small open circles). The experiment was repeated three times independently, with similar results. **(C)** Representative image of ten-day-old seedlings of WT and *miPEP858^CR^* lines (L-1, L-3, L-5) grown on half-strength MS medium supplemented with water (control) and 0.25 µM 1F, 2F, and 3F. Scale bar, 1 cm. **(D)** Root lengths of ten-day-old seedlings of WT and *miPEP858^CR^* lines (L-1, L-3, L-5) grown on half-strength MS medium supplemented with water (control) and 0.25 µM 1F, 2F, and 3F (n= 30 independent seedlings, indicated by the small open circles). The experiment was repeated three times independently, with similar results. **(E)** Representative image of 30-day-old WT and *miPEP858^CR^* lines (L-1 and L-3) supplemented with water (control) and 0.25 µM 2H, 3F and miPEP858a. **(F)** Bolting time of WT and *miPEP858^CR^*lines (L-1 and L-3) grown under standard light conditions (16h light/ 8 h dark) supplemented with water (control) and 0.25 µM 2H, 3F and miPEP858a. n= 15 (the small open circles represent the individual values). **(G)** Rosette diameter of 30-day-old WT and *miPEP858^CR^* lines (L-1 and L-3) supplemented with water (control) and 0.25 µM of 2H, 3F and miPEP858a. The statistical analysis was performed using two-tailed Student’s t-tests. The data are plotted as means ± s.d. The error bars represent standard deviations. The asterisks indicate significant differences; *P < 0.1; **P < 0.01; ***P < 0.001.

The effect of truncated peptides on root length and development of *miPEP858^CR^* further prompted us to demonstrate the effect of these functional truncated peptides on the mature WT and *miPEP858^CR^*plants (Figure 3E). Exogeneous application of truncated peptides suggested the early bolting phenotype in plants treated with truncated peptides than the plants treated with water (Figure 3F). Since the overall growth was enhanced after external application of truncated peptides, we also analyse the rosette diameter of the Col-0 and *miPEP858^CR^* plants treated with the functional truncated peptides. Analysis suggested the overall increase in rosette diameter of peptide-treated WT as well as *miPEP858^CR^* plants (Figure 3G). Together, these results suggest that the exogenous application of functional truncated peptides has the potential to complement *miPEP858^CR^* at the phenotypic level.

To investigate the molecular impact of functional truncated peptides on *miPEP^CR^* lines, we analyse the expression of miR858 and its target and associated genes in *miPEP858^CR^* seedlings supplemented with 2H and 3F peptides. The expression analysis suggests the increase in expression of miR858 and decrease in expression of its target and associated genes (MYB12, CHS) in *miPEP858^CR^*seedlings when supplemented with the 2H and 3F truncated peptides (Figure 4A). We also studied the MYB12 and CHS protein accumulation in response to exogeneous supplementation of functional truncated peptides and it suggested the lesser protein accumulation in *miPEP858^CR^* seedlings treated with 2H and 3F peptide compare to control (water), *miPEP858^CR^* and WT seedlings (Figure 4B). Lignin staining of cross-sections of 35-day old mature stem under the microscope revealed lesser lignification in the vascular tissues and interfascicular fibres of lignin in *miPEP858^CR^*control plants compared to the plants treated with functional 2H and 3F truncated peptides (Figure 4C). Measurement of length of vascular bundle and interfascicular fibre suggested an increase in length of vascular bundle and interfascicular bundle when treated with functional truncated peptide compare to control plants (Figure 4D and 4E). Estimation of total flavonols and anthocyanins content also suggested that the exogenous supplementation of functional truncated peptide led to a decrease in flavonols and anthocyanins content in *miPEP858^CR^* seedlings compared to water-treated *miPEP858^CR^* seedlings (Figure 4F, and 4G). These results suggest that the exogenous application of C-terminal peptide has the potential to complement the function of miR858 in *miPEP858^CR^*lines and also lead to modulation in gene expression associated with the phenylpropanoid pathway.

**Figure 4.**
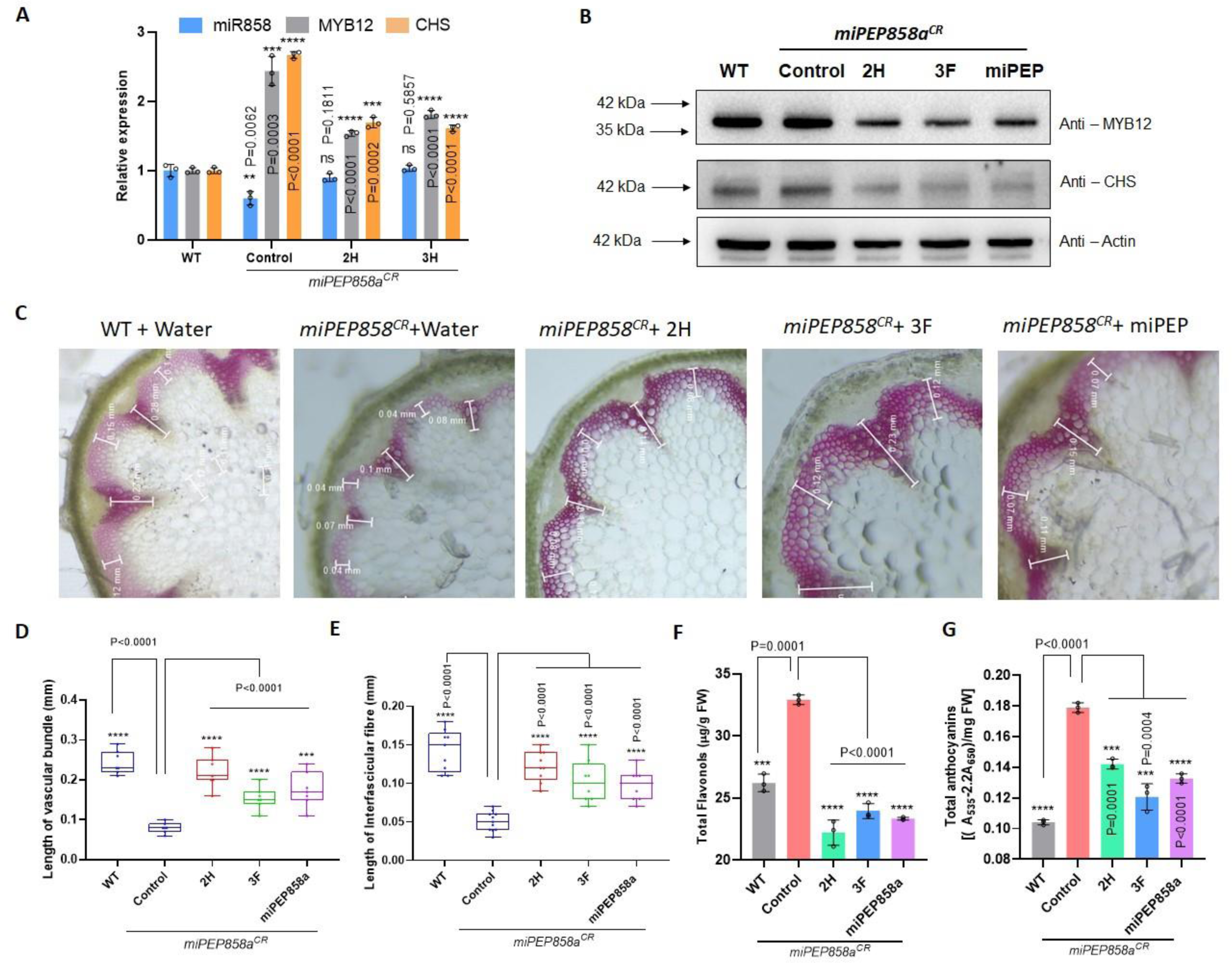
Modulation of expression of miR858 and target genes and metabolite accumulation on supplementation with truncated peptide. **(A)** Quantification of *miR858a* and its target genes *MYB12* and *CHS* in five-day-old seedlings of WT grown on half-strength MS medium supplemented with water and *miPEP858^CR^* lines supplemented with water (control) and 0.25 µM of truncated peptides 2H and 3F. **(B)** Western blot analysis of MYB12 and CHS protein in five-day-old WT seedlings grown on half-strength MS medium supplemented with water and *miPEP858^CR^* lines supplemented with water (control) and 0.25 µM of 2H, 3F and miPEP858a. Actin was used as the loading control. **(C)** Quantification of lignin in 35-day-old stems of WT plants supplemented with water and *miPEP858^CR^* lines supplemented with water (control) and 0.25 µM of 2H, 3F and miPEP858a. The experiments were repeated three times independently, with similar results. **(D)** Lengths of vascular bundles of 35-day-old stems of WT plants supplemented with water and *miPEP858^CR^*lines supplemented with water (control) and 0.25 µM of 2H, 3F and miPEP858a (n=15). **(E)** Lengths of interfascicular fibres of 35-day-old stems of WT plants supplemented with water and *miPEP858^CR^* lines supplemented with water (control) and 0.25 µM of 2H, 3F and miPEP858a (n=15). **(F)** Quantification of total flavonol in ten-day-old WT seedling grown on half-strength MS medium supplemented with water and *miPEP858^CR^* lines supplemented with water (control) and 0.25 µM of 0.25 µM of 2H, 3F and miPEP858a. FW, fresh weight. **(G)** Quantification of anthocyanin in ten-day-old WT seedlings grown on half-strength MS medium supplemented with water and *miPEP858^CR^* lines supplemented with water (control) and 0.25 µM of 0.25 µM of 2H, 3F and miPEP858a. FW, fresh weight.

### C-terminal truncated peptide regulates PSK4

miPEP858 is recognized for upregulating miR858 transcription, and the exogenous application of miPEP858 to promoter lines enhances the expression of the reporter gene (Sharma et al., 2020). Expanding our investigation, we examined the impact of truncated peptides on miPEP858 promoter lines. GUS histochemical staining of miPEP858 promoter lines (Pro:ATG1::GUS and Pro:ORF1::GUS) supplemented with truncated peptides indicated increased GUS expression in promoter:reporter lines treated with functional 2H and 3F peptides. Conversely, other truncated peptides showed no significant change in histochemical GUS staining (Figure 5A and 5C). These findings were further corroborated by expression analysis of the GUS gene, revealing increased expression in ATG1 and ORF promoter seedlings treated with 2H and 3F truncated peptides (Figure 5B and 5D).

It has been reported that synthetic miPEP858 enhances the expression of the PSK4 transcript (Badola et al., 2022). To assess the impact of truncated peptides on PSK4 expression, WT seedlings were treated with truncated peptides, and the analysis revealed an increased PSK4 transcript only in response to the supplementation of 2H and 3F functional truncated peptides. In contrast, other truncated peptides failed to modulate the expression of the PSK4 gene (Figure 5E). The analysis of auxin-related genes indicated a significant increase in their expression in seedlings treated with 2H and 3F peptides. Conversely, non-functional truncated peptides did not induce changes in auxin-related gene expression (Figure 5F, Supplemental Figure 9A). Considering that PSK4 overexpression lines lead to increased expression of growth-related expansins (Badola et al., 2022 30), we analyzed the expression of these expansins in truncated peptide-treated seedlings. The analysis suggested an increase in the expression of growth-related expansins in WT seedlings supplemented with 2H and 3F functional truncated peptides (Figure 5G, Supplemental Figure 9B). Exogenous application of synthetic miPEP858 increases GUS expression in PSK4 promoter seedlings; therefore, we performed histochemical GUS staining and expression analysis of the GUS gene in PSK4 promoter lines. Both histochemical GUS staining and expression analysis indicated an increase in GUS gene expression in PSK4 promoter seedlings supplemented with 2H and 3F functional truncated peptides (Figure 5H and 5I). Overall, these results suggest the potential of truncated functional peptides to enhance miPEP858 and PSK4 expression, along with various growth-related genes (auxin-and growth-related expansins), thereby influencing plant growth and development.

## Discussion

Small signaling peptides fall into two categories: secreted and non-secreted peptides, serving as vital signaling molecules for cell-to-cell communication. Many of these peptides are translated as precursor proteins, giving rise to mature peptides that function in diverse developmental and physiological processes (Mastubayashi and Sakagami 2006). Small peptides, such as PSK, CLE, IDA, CEP, RGF, and others, play crucial roles in root development, stomatal conductance, abscission processes, stem cell maintenance, root elongation, and stress response (Qian et al., 2018, Aalen et al., 2013, Olsoon et al., 2019, Takahashi et al., 2018, Datta et al., 2024). Typically, small signaling peptides, whether secreted or non-secreted, undergo proteolytic processing to generate a mature signal peptide, which is then post-translationally modified. The N-terminal segment processes most small signaling peptides, with their functional C-terminal regions responsible for their biological functions. In the case of non-secreted peptides, these molecules are released directly from cells, executing their functions without the need for processing and maturation (Mastubayashi 2011). One example is AtPEP1, a small peptide that triggers signals for the innate immune response against pathogen attacks (Yamaguchi et al., 2006). Similarly, AtPEP3, another peptide from the same family, is involved in salinity stress tolerance. Its C-terminal peptide is activated in response to salinity, inhibiting salinity-induced chlorophyll bleaching during stress conditions (Nakaminami 2018). In summary, small peptides play significant roles in plant developmental processes and responses to stress.

miRNA858 has the potential to regulate the phenylpropanoid pathway by targeting MYB transcription factors (MYB111, MYB11, MYB12) (Sharma et al., 2016). The primary sequence of miR858 encodes a regulatory peptide, miPEP858. Exogenous application of miPEP858a not only affects plant growth and development but also modulates secondary plant products (Sharma et al., 2020). While miR858 plays a role in the phenylpropanoid pathway, major growth changes depend on downstream molecular components, including the small signaling peptide PSK4. The crosstalk between AtPSK4 and miR858/MYB3 is crucial for plant growth and development, underscoring the significance of these small peptides. Despite this, exogenous application of miPEP858a significantly induces the expression of the reporter gene GUS in Pro:PSK4 lines, highlighting the importance of miPEP858 in controlling the PSK4 peptide (Badola et al., 2022). In summary, small peptides effectively regulate various developmental and physiological processes. In our previous study (Sharma et al., 2020), CRISPR/Cas9-based mutants were developed, resulting in edited lines with altered growth patterns and secondary metabolite levels. Sequence analysis of miPEP858 edited lines revealed a truncated amino acid sequence at the C-terminus while maintaining the intact N-terminus, emphasizing the essential role of C-terminal amino acids in the functionality of miPEP858a.

While some miPEPs play known roles in various developmental processes, their exogenous application for controlling desired traits in plants can be prohibitively expensive. Custom-made synthetic peptides are costly, and achieving an optimal concentration for a functional phenotype often necessitates a higher peptide amount for exogenous treatment. To address the cost concern, a minimal 13-amino acid fragment of miPEP858 has been identified, reducing synthesis expenses while providing a similar effect as the full-length 44-amino acid miPEP858. This minimal fragment has been demonstrated to exhibit a comparable phenotype and modulation in the accumulation of flavonols/anthocyanins, similar to full-length miPEP858. To identify functional truncated peptides, several designs were explored from the N and C-terminus ends of miPEP858, including 1^st^ half, 2^nd^ half, 1^st^ fragment, 2^nd^ fragment, and 3^rd^ fragment (Figure 1D). Analysis revealed that peptides from the C-terminus end (2H and 3F) provided a response similar to miPEP858. Hence, it was hypothesized that 2H and 3F fragments could serve as functional truncated peptides, influencing the phenotype and modulating the expression of flavonoid biosynthesis pathway genes. Expression analysis of miR858 indicated an increase in expression in response to 2H and 3F. Additionally, significant changes in the expression of MYB12 and CHS were observed in response to functional truncated peptides compared to the other three non-functional peptides, with lower protein accumulation in the case of functional truncated peptides (Figure 2A – C and Supplemental Figure 2).

This research underscores the significance of the C-terminus end of miPEP858, housing crucial functional amino acids essential for the phenotype and modulation of secondary plant products, particularly flavonols and anthocyanins. Clathrin-mediated endocytosis stands out as a key internalization route in plants (Dhonukshe 2007, Kitakura 2011), with studies suggesting the internalization of miPEP165 and miPEP858 through this mechanism. To investigate the uptake of truncated peptides, FAM-labelled 3F and 2H peptides were incubated, revealing a mode of uptake similar to miPEP165 and miPEP858 inside the plant cell (Ormancey et al., 2020, Badola et al., 2022). Earlier reports indicate that miPEP can enhance and regulate the activity of its own promoter (Waterhouse and Hellens 2015). Promoter reporter studies also suggested that the exogenous application of miPEP858 on PromiR858a:ATG1::GUS and PromiR858a:ORF1::GUS enhanced GUS staining (Sharma et al., 2020). Building on this, we hypothesized that if the C-terminus is functional in modulating the phenotype and expression of key genes, it should increase GUS expression. As anticipated, enhanced GUS staining was observed when promoter seedlings were supplemented with 2H and 3F along with miPEP858 (Figure 5A - D), suggesting that the C-terminus functions similarly to full-length miPEP858.

Our earlier study indicates that full-length miPEP858a regulates plant growth and development by modulating the expression of PSK4, enhancing the expression of growth-related auxin and expansin genes (Badola et al., 2022). To further validate the function of truncated peptides, the expression of PSK4 was analyzed in Arabidopsis seedlings supplemented with all truncated peptides. The analysis suggested increased expression in response to supplementation with peptides derived from the C-terminal end. The effect of functional truncated peptides was also analyzed on Pro: PSK4 seedlings supplemented with functional truncated peptides, showing a significant increase similar to full-length miPEP858. Like miPEP858, C-terminus truncated peptides exhibited a significant increase in the expression of auxin and expansins, key genes required for growth and development (Figure 5F and 5G).

**Figure 5.**
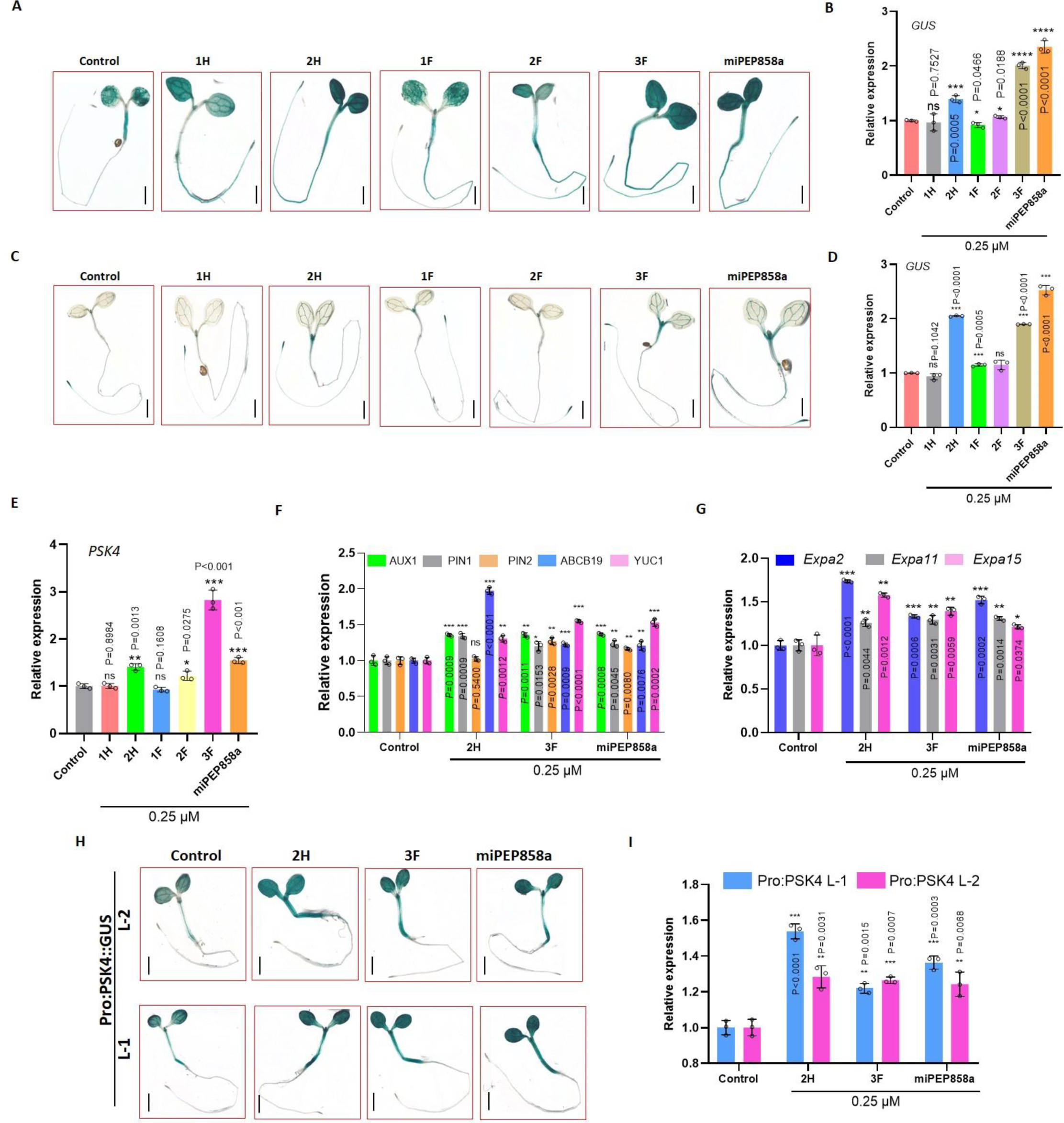
Effect of truncated peptide on down-stream processes to regulate growth and development. **(A)** Histochemical staining showing GUS activity in five-day-old transgenic seedlings of Pro: ATG1:: GUS grown on half-strength MS medium supplemented with water (control) and 0.25 µM of 1H, 2H, 1F, 2F, 3F and miPEP858a. Scale bars, 1,000 µm. The experiment was repeated three times independently (n=15 seedlings), with similar results. **(B)** Relative expression of GUS in Pro: ATG1:: GUS transgenic seedlings grown on half-strength MS medium supplemented with water (control) and 0.25 µM of 1H, 2H, 1F, 2F, 3F and miPEP858a. **(C)** Histochemical staining showing GUS activity in five-day-old transgenic seedlings of Pro: ORF1::GUS grown on half-strength MS medium supplemented with water (control) and 0.25 µM of 1H, 2H, 1F, 2F, 3F and miPEP858a. Scale bars, 1,000 µm. The experiment was repeated three times independently (n=15 seedlings), with similar results. **(D)** Relative expression of GUS in Pro: ORF1:: GUS transgenic seedlings grown on half-strength MS medium supplemented with water (control) and 0.25 µM of 1H, 2H, 1F, 2F, 3F and miPEP858a. **(E)** Quantification of *AtPSK4* in five-day-old WT seedlings grown on half-strength MS medium supplemented with water (control) and 0.25 µM of 1H, 2H, 1F, 2F,3F and miPEP858a exogenously. **(F)** Relative expression of auxin genes (*AUX1*, *PIN1*, *PIN2*, *ABCB19* and *YUC1*) in five-day-old WT seedlings grown on half-strength MS medium supplemented with water (control) and 0.25 µM of functional truncated peptides 2H, 3F and miPEP858a. **(G)** Relative expression of *EXPA2*, *EXPA11* and *EXPA15* gene expression in five-day-old WT seedlings grown on half-strength MS medium supplemented with water (control) and 0.25 µM of functional truncated peptides 2H, 3F and miPEP858a **(H)** Histochemical staining showing GUS activity in five-day-old transgenic seedlings of Pro: PSK4:: GUS lines (L-1 and L-2) grown on half-strength MS medium supplemented with water (control) and 0.25 µM of functional truncated peptides 2H,3F and miPEP858a. Scale bars, 1,000 µm. The experiment was repeated three times independently (n=15 seedlings), with similar results. **(I)** Relative expression of GUS in seedlings of Pro: PSK4:: GUS transgenic lines (L-1 and L-2) grown on half-strength MS medium supplemented with water (control) and 0.25 µM of functional truncated peptides 2H, 3F and miPEP858a.

Based on our study, we propose a model illustrating the role of the C-terminal region of miPEP858, functioning almost similarly to full-length miPEP858 in modulating phenylpropanoid pathway genes as well as plant growth and development. Consequently, we conclude that C-terminal peptides offer a cost-effective approach to enhancing agronomic traits without the need for cumbersome biotechnological methods (Figure 6). Ultimately, our results underscore the importance of identifying minimal functional fragments as an alternative for exogenous use, providing not only cost-effectiveness but also the potential to enhance agronomic traits. This research establishes a groundwork for identifying more functional peptides across various species and using them exogenously to improve desired agronomic traits. Synthetic peptides can effectively be employed to enhance agronomic traits in crop plants, contributing to advancements in food safety and security in the future.

**Figure 6.**
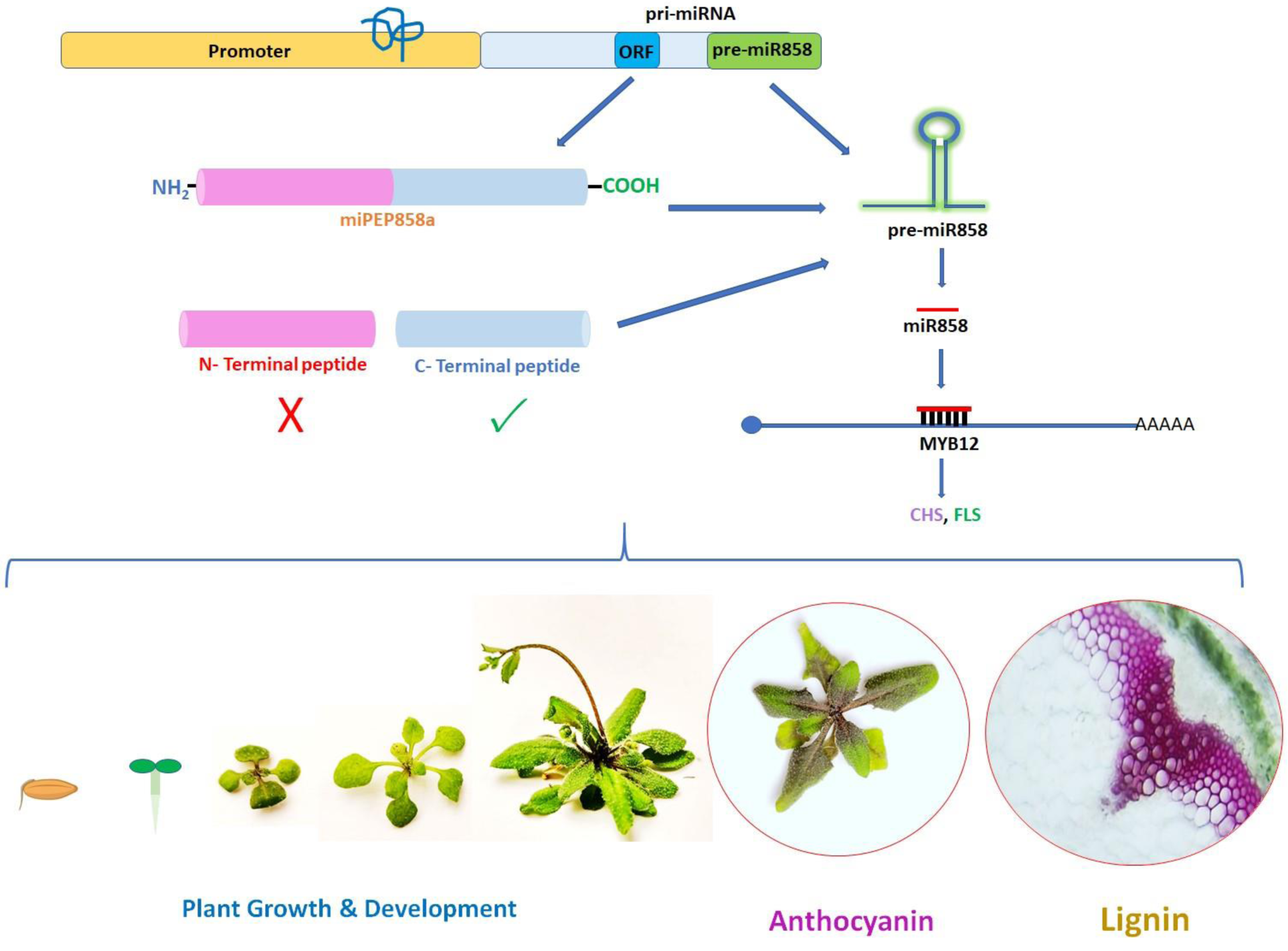
Model of the study showing the C-terminus region of miPEP858a can enhance the expression of precursor and mature miR858a similar to full-length miPEP858a providing evidence regarding the importance of amino acids present at the C-terminus end. The C-terminus peptides have the potential to regulate phenylpropanoid pathway genes, and metabolites along with plant growth and development.

## Materials and methods

### Plant material and growth conditions

*Arabidopsis thaliana* (Col-0) was used as the wild type (WT) throughout the study. Firstly, seeds were surface sterilised and then placed on half-strength MS medium (Hi-Media) containing 1.5% sucrose, pH ∼5.72. After 2 days of stratification in the dark at 4^°^C, the plates were transferred to a Growth Chamber (Percival) set at a 16-h-light/8-h-dark photoperiod cycle, 180 µmolm^−2^ s^−1^ light intensity, 22-24°C temperature and 60–70% relative humidity. Vertically grown 10-day-old seedlings were used for the root length measurements. The *miR858^CR^*, *miPEP858^CR^*, ATG-GUS, ORF-GUS, and Pro:PSK4 lines were previously generated by our group (14, 30) and *chc1* AT3G11130 SALK_112213, *chc1.2*, SALK_103252, *chc2* AT3G08530 SALK_042321, and *chc2.1* SALK_028826 mutants were obtained from ABRC used in this study for analysis. To study the effect of all truncated peptides, Col-0 seedlings grown on media supplemented with water (control) and all the truncated peptides along with miPEP858 as a positive control throughout the study.

### Synthetic peptide assay

The synthetic peptides having purity>95% were synthesized through Link Biotech http://www.linkbiotech.com/. All peptides were dissolved in water (stock concentration, 5 mM). The seedlings were treated with different concentrations, ranging from 0.1 to 0.5 μM peptide, diluted in the agar medium or in water for spraying on the mature plants. For mature plants grown in soilrite, 14-and 21-day-old plants were treated with peptide (0.5 μM) diluted in water. Following peptides were used for the study:

> **miPEP858a:** MGGIESLLFTIVRDIGRYGTVCVVYNIKCVYTTRTKASTRTSHP
>
> **1st half [1H]:** MGGIESLLFTIVRDIGRYGTVC
>
> **2nd half [2H]:** VVYNIKCVYTTRTKASTRTSHP
>
> **1st fragment [1F]:** MGGIESLLFTIVRDI
>
> **2^nd^ fragment [2F]:** GRYGTVCVVYNIKCV
>
> **3^rd^ fragment [3F]:** YTTRTKASTRTSHP

### miPEP858a structure prediction (folding) and molecular docking

The miRNA858a location in Arabidopsis was retrieved from the miRBase database (https://www.mirbase.org/browse/results/?organism=ath). The promoter and miRNA precursor regions were extracted based on the provided locations for the miRNA. Functional open reading frames (miPEP858a) were then translated for their protein-coding strands. Subsequently, the interaction between miPEP858a and the promoter sequence of miPEP858a was investigated. The identified miRNA peptide was structurally predicted using the PEP-FOLD server (https://bioserv.rpbs.univ-paris-diderot.fr/services/PEP-FOLD/). The three-dimensional structure of the promoter region was predicted through the make-na server (https://web.archive.org/web/20170430174556/http:/structure.usc.edu/make-na/server.html). Docking analysis was performed to examine the interaction between the promoter (DNA) and miPEP858a using the NPDock server (https://genesilico.pl/NPDock/).

### Chromatin immunoprecipitation

ChIP assays were conducted using 3-week-old seedlings. Six grams of seedling tissues were weighed and cross-linked in 37 ml of 1% formaldehyde mixed with 50 ml extraction buffer 1 [0.4 M sucrose, 10 mM Tris-HCl, pH 8, 5 mM beta-mercaptoethanol, 0.1 mM phenylmethylsulfonyl fluoride (PMSF), and Protease inhibitor cocktail; sigma]. Cross-linking was performed under vacuum for 10 minutes, followed by the addition of 2.5 ml of 2M glycine to stop the process. After rinsing the seedlings twice with cold autoclaved water, excess water was removed, and the samples were stored at −80°C. The next day, chromatin isolation was initiated by grinding the tissue to a fine powder using a pre-cooled mortar and pestle. The powder was resuspended in 20 ml of Extraction buffer I, filtered through a cell strainer, and centrifuged. The pellet was successively resuspended in Extraction buffers 2 [0.25 M sucrose, 10 mM Tris-HCl, pH 8, 10 mM MgCl2, 1% Triton X-100, 5 mM beta-mercaptoethanol, 0.1 mM phenylmethylsulfonyl fluoride (PMSF), and Protease inhibitor cocktail; sigma] and 3 [1.7 M sucrose, 10 mM Tris-HCl, pH 8, 0.15% Triton X-100, 2mM MgCl2, 5 mM beta-mercaptoethanol, 0.1 mM phenylmethylsulfonyl fluoride (PMSF), and Protease inhibitor cocktail; sigma], followed by centrifugation. The crude pellet was suspended in nuclei lysis buffer, sonicated to achieve an average fragment size of 0.3 to 1.0 kb, and checked for sonication efficiency. Magnetic beads (A/G) were dissolved in ChIP dilution buffer and used to pre-clear the sonicated chromatin. The sonicated chromatin was then incubated overnight with miPEP858-specific antibodies. After washing, the elution buffer was added to the beads, and the solution underwent an elution process. The eluate was subjected to a reverse cross-linking step, and DNA was extracted, precipitated, washed, and resuspended in TE buffer. A small aliquot of the extracted DNA was used for qRT-PCR analysis.

### Yeast one-hybrid assay

For the yeast one-hybrid assay, the coding sequence of miPEP858a was amplified and cloned into the pGADT7AD vector to generate a fusion with the Activation domain. The promoter sequence of miPEP858a (approximately 1500 bp upstream of the start codon) was amplified and cloned into the pAbAi vector to create the DNA bait sequence. To investigate the interaction between the miPEP858a peptide and the miPEP858a promoter, the pAbAi vector containing the miPEP858a promoter was linearized, transformed into the Y1HGold yeast strain, and allowed to grow on SD/-Ura media. Positive transformant colonies were confirmed by PCR, followed by selection on Aureobasidin A (AbA). In the next step, the pGADT7 vector containing the miPEP858a coding sequence was transformed into the Y1HGold [Bait] strain and allowed to grow on SD/-Leu/þAbA media.

### Confocal microscopy imaging

For peptide uptake assays in Arabidopsis roots, fluorescent carboxyfluorescein (5-FAM)-labeled peptides were purchased from Link Biotech (95%–98% purity). Three-day-old Arabidopsis WT and *chc* mutant seedlings were incubated with 5-FAM-labelled peptides (50 µM) in MG buffer (10-mM MgCl_2_ buffer, pH 5.8) at 22°C for 12 hours. After incubation with labelled peptide, the seedlings were washed 3 times by gentle shaking for 5 min in MG buffer, and the roots were analyzed under confocal microscope (Zeiss LSM710, Zeiss LSM Image Examiner Version 4.2.0.121, CarlZeiss) at an excitation 495 nm/emission 545 nm and solid-state laser (10% of 10 mW) for both FAM and PI. Roots were observed under Leica microscope (LAS version 4.12.0, Leica Microsystems.

### Gene expression analysis

The gene expression was analysed through qRT-PCR (quantitative Real-time PCR). Total RNA was extracted from different samples, treated with DNAase (Thermo), and (1µg) reverse transcribed using the Revert AidH Minus First Strand cDNA Synthesis Kit (Thermo) as per the manufacturer’s instructions. The cDNA was diluted 20 times with nuclease-free water, and 2 µl was used as a template for qRT-PCR performed using Fast SYBR Green Mix (Applied Biosystems) in a Fast 7500 Thermal Cycler instrument (Applied Biosystems). The expression was normalised using Tubulin and analysed through the comparative ΔΔ^CT^ method (Schmittgen and Livak 2008). For the expression analysis of mature miR858a, TaqMan PCR assays were used following the manufacturer’s protocol (Applied Biosystems). Small nuclear RNA (snoR41Y) was used as a normalization control. The primer sequences used in the study are listed in **Supplementary Table S1**.

### Histochemical GUS staining

The histochemical GUS staining was performed using a previously described method (Jeferson et 1989). Promoter:reporter seedlings of the Pro:miPEP858:ATG:GUS, Pro:miPEP858:ORF:GUS and ProPSK4:GUS transgenic lines were dipped in a solution containing 100-mM sodium phosphate buffer (pH 7.2), 10-mM EDTA, 0.1% Triton X-100, 2-mM potassium ferricyanide, 2-mM potassium ferrocyanide, and 1-mg mL^−1^ 5-bromo-4-chloro-3-indolyl-b-D-glucuronide (X-GluC) at 37°C for 4-6 hours. After staining, chlorophyll was removed by incubation and multiple washes using 70% ethanol. The seedlings were observed under a Leica microscope (LAS version 4.12.0; Leica Microsystems, Wetzlar, Germany) for the GUS staining.

### Total flavonol and anthocyanin quantification

Five-day-old seedlings were extracted in 1 ml of 80% methanol at 4°C for 2h with shaking. The mixture was centrifuged at 12,000 g for 12 min. The supernatants (0.5 ml) were taken to 2 ml with methanol and subsequently mixed with 0.1 ml of aluminium chloride (10% water solution), 0.1 ml of potassium acetate (1 M) and 2.8 ml of MQ water. After 30 min of incubation, the absorbance was taken at 415 nm. The calibration curve was developed using rutin as the standard. The total flavonol content was calculated as the equivalents of rutin used as the standard (Loyala et al., 2016). Total anthocyanin content was quantified according to the previously described method (Li et al, 2016). Briefly, the five-day-old seedlings (300 mg) were crushed in liquid N_2_ and transferred in the extraction solution (Isopropanol: HCl: H_2_O::18:1:81). The samples were then heated at 95°C for 3 min, followed by incubation at room temperature in the dark for 2h. After centrifugation, the absorbance of the supernatants was measured at A_535_ and A_650_. Total anthocyanin content was calculated as (A_535_−2.2A_650_)/g FW.

### Extraction and quantification of flavonols

For the extraction of flavonols, the seedlings (300 mg) were ground in liquid nitrogen and extracted in 80% methanol overnight at room temperature. Total extracts were hydrolysed in an equal amount of 6N HCl at 70 °C for 40 min. This was further proceeded by the addition of an equal amount of methanol to prevent the precipitation of the aglycones. The extracts were filtered through 0.2 µm filters (Millipore) before the metabolite analysis using HPLC. All the samples were analysed by HPLC-PDA with a Waters 1525 Binary HPLC Pump system comprising PDA detector following the method developed by Niranjan et al (Niranjan et al., 2011). Breeze 2 software (Waters) was used for the quantification of various metabolites through HPLC.

### Lignin staining

To visualize lignified cells in stems, hand cut sections from 35-day-old mature plants were stained using phloroglucinol (Sigma-Aldrich) for 1 min and visualized on a Leica DM2500 microscope.

### Total protein extraction and western blot analysis

Total protein extraction and western blot analysis were essentially carried out as per the methods described in Sharma et al. (Sharma et al., 2020). The western blot images were captured using Image Lab version 5.2.1 build 11 (Bio-Rad Laboratories). The commercial antibodies used in the analysis were Anti-Actin Antibody (A0480, Sigma Aldrich), anti-CHS (AS122615, Agrisera). The polyclonal antibodies were generated in rabbit against a peptide (CEGSDNNLWHEKENP) in the MYB12 protein sequence and were affinity purified against the peptide (Eurogenetec).

### Statistical analysis

The statistical tests and n numbers, including sample sizes or biological replications, are described in the figure legends. All the statistical analyses were performed using two-tailed Student’s t-tests using GraphPad Prism version 9.0 software. All the experiments were repeated at least three times independently, with similar results.

## Supplemental data

**Supplemental Figure 1.** Binding of miPEP858a on its promoter.

**Supplemental Figure 2.** Effect of functional and non-functional peptide on phenylpropanoid pathway genes.

**Supplemental Figure 3.** Quantification of flavonols.

**Supplemental Figure 4.** Internalization of FAM labelled truncated peptide (2H) in clathrin mutants.

**Supplemental Figure 5**: Internalization of FAM labelled truncated peptide (3F) in clathrin mutants.

**Supplemental Figure 6.** Effect of function truncated peptide on growth of clathrin mutants.

**Supplemental Figure 7.** Effect of non-function truncated peptide on growth of clathrin mutants.

**Supplemental Figure 8.** Exogenous application of functional truncated peptide fail to complement *miR858^CR^* lines.

**Supplemental Figure 9.** Effect of non-functional peptide on Auxin and Expansin genes.

**Supplemental Table 1.** List of primers used in the study

## Acknowledgement

This research was supported by the Council of Scientific and Industrial Research (CSIR), New Delhi, in the form of NCP project no. MLP006. P.K.T. also acknowledges Science and Engineering Research Board, New Delhi for JC Bose National Fellowship (JCB/2021/000036). H.G., A.A., acknowledges the Council of Scientific and Industrial Research and Department of Biotechnology New Delhi, for a Senior and Junior Research Fellowship. A.S. acknowledge the Department of Science and Technology for the DST-Inspire Faculty Project. Authors also acknowledge Dr. Manju Singh from Central Instrumentation Facility, CSIR-CIMAP for phytochemical analysis, and Dr. Sanchita, CSIR-CIMAP and Dr. Mehar Hasan Asif from CSIR-NBRI for bioinformatic analysis.

## Data availability

All data generated or analyzed during this study are included in this published article (and its supplementary information files).

